# Scalable and model-free detection of spatial patterns and colocalization

**DOI:** 10.1101/2022.04.20.488961

**Authors:** Qi Liu, Chih-Yuan Hsu, Yu Shyr

## Abstract

The expeditious growth in spatial omics technologies enable profiling genome-wide molecular events at molecular and single-cell resolution, highlighting a need for fast and reliable methods to characterize spatial patterns. We developed SpaGene, a model-free method to discover any spatial patterns rapidly in large scale spatial omics studies. Analyzing simulation and a variety of spatial resolved transcriptomics data demonstrated that SpaGene is more powerful and scalable than existing methods. Spatial expression patterns by SpaGene reconstructed unobserved tissue structures. SpaGene also successfully discovered ligand-receptor interactions through their colocalization.

## INTRODUCTION

Spatial omics technologies map out organizational structures of cells along with their genomics, transcriptomics, proteomics and epigenomics profiles, providing powerful tools for deciphering mechanisms of functional and spatial arrangements in normal development and disease pathology (Larsson et al. 2021; Longo et al. 2021; Marx 2021; Deng et al. 2022; Dhainaut et al. 2022; Ratz et al. 2022; Zhao et al. 2022). The collection of available approaches provides a wide spectrum of throughput and spatial resolution. Imaging-based approaches generally target pre-selected RNA or proteins at molecular and single cell resolution, while sequencing-based approaches allow genome-wide profiling with limited spatial resolution (Lewis et al. 2021; Zhuang 2021). Recent advances in those approaches move the field rapidly into the direction achieving both high throughput and spatial resolution, presenting a significant computational challenge for scalable and robust methods to derive biological insights in the spatial context (Atta and Fan 2021).

One essential step in spatial omics analysis is to characterize spatial expression patterns and colocalization. Several methods have been developed to identify spatially variable genes (Edsgard et al. 2018; Svensson et al. 2018; Sun et al. 2020a; Anderson and Lundeberg 2021; Miller et al. 2021; Zhu et al. 2021). Trendsceek uses permutation test to detect significant dependency between the spatial distribution of points and their expression levels based on marked point processes (Edsgard et al. 2018). Sepal ranks spatially variable genes by the diffusion time with the rational that genes with spatial patterns require more time to reach a homogenous state than those with random spatial distributions (Anderson and Lundeberg 2021). SpatialDE and SPARK both utilize Gaussian process regression as the underlying data generative model for spatial covariance structures. SpatialDE decomposes expression variability into spatial variance and noise, and estimates statistical significance by comparing the likelihoods with and without a spatial component (Svensson et al. 2018). SPARK extends SpatialDE via generalized linear spatial error models, with the ability to directly model raw counts and adjust for covariates (Sun et al. 2020a). SPARK-X examines the similarity of expression covariance matrix and distance covariance matrix and tests whether they are more similar than expected by chance (Zhu et al. 2021). The statistical power of such methods highly depends on spatial covariance models, i.e, how well they match true underlying expression patterns. Although multiple kernels, including Gaussian, linear and periodic kernels with different smoothness parameters, are considered to ensure identification of various spatial patterns, statistical power will be compromised substantially for identifying spatial patterns poorly modelled by those predefined kernel functions. Furthermore, spatial covariance models are built upon cellular distances, which would confound true expression variances with those driven by variances in cellular densities. To take non-uniform cellular densities into consideration, MERINGUE calculates spatial autocorrelation and cross-correlation based on spatial neighborhood graphs to identify spatially variable genes and gene interactions (Miller et al. 2021). Above all, even equipped with computationally efficient algorithms, it would still take days to months for most methods to analyze large-scale spatial data with genome-wide profiling in tens of thousands of locations (Zhu et al. 2021), resulting in a high demand for scalable and robust methods for characterizing spatial expression patterns.

Here we developed SpaGene, a scalable and model-free method for detecting spatial patterns. SpaGene is built upon a simple intuition that spatially variable genes have uneven spatial distribution, meaning that cells/spots with high expression tend to be more spatially connected than random. SpaGene is one of the most computationally efficient methods, which only takes seconds to minutes for analyzing large-scale spatial omics data. Independent of spatial covariance models and cellular densities, SpaGene demonstrated the power to identify any spatial patterns in the simulation and a variety of spatial transcriptomics datasets. Spatial expression patterns by SpaGene reconstructed unobserved tissue structures. Extended to identify spatial colocalization, SpaGene successfully discovered cell-cell communications mediated by ligand-receptor interactions.

## RESULTS

### Simulation

A schematic diagram of SpaGene is shown in Fig. 1A, with details in the Methods section. We first applied SpaGene on two simulation datasets. One simulation was generated from negative binomial distributions following SPARK-X (Zhu et al. 2021), the other was sampled from real data following Trendsceek (Edsgard et al. 2018). Cells/spots with higher expression (spiked cells) were located in one of those five patterns, hotspot, streak, circularity, bi-quarter circularity, and Purkinje layer in mouse cerebellum (Fig. 1B). The distinctness of the pattern was determined by effect sizes, which were controlled by the fold change (FC) of expression in spiked cells compared to the background. The pattern size was determined by the percentage of spiked cells. Higher effect sizes and larger pattern sizes generated more distinct and bigger patterns, which were easier to be identified. Among the simulated genes, 500 genes display spatial patterns (details in the Methods section). The area under the curve (AUC) was used to measure the ability to distinguish between spatially and non-spatially variable genes.

**Fig. 1.**
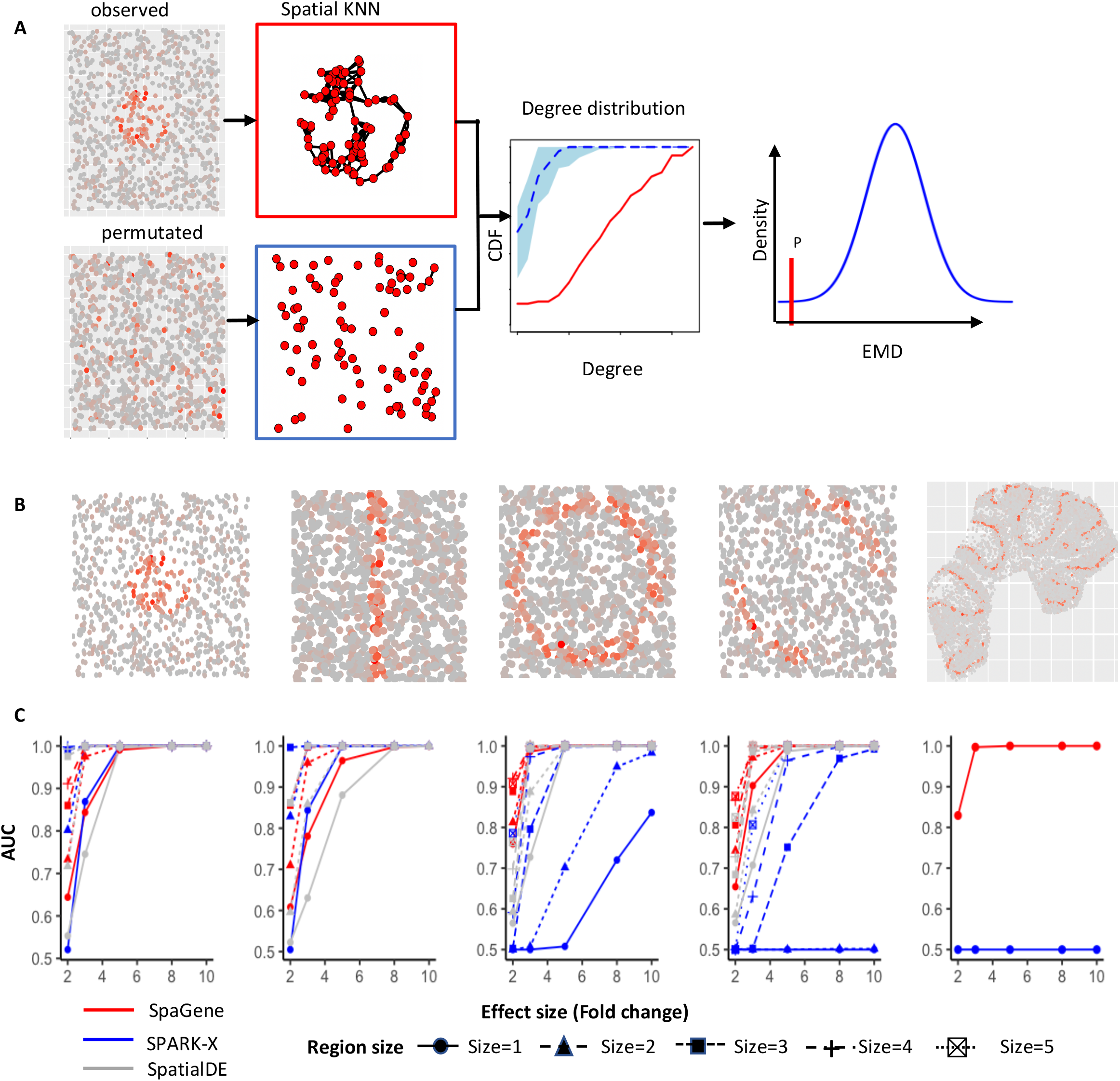
Schematic of SpaGene and simulation results. A) Schematic of SpaGene; B) Visualization of five spatial patterns; C) AUC plots of SpaGene (red), SpatialDE (gray) and SPARK-X (blue) in simulated datasets with different effect sizes (x axis) and pattern sizes (point shapes) and 10,000 genes and 1,000 cells/locations. Simulated data were generated from negative binomial distributions.

We compared SpaGene with SpatialDE and SPARK-X. SpatailDE and SPARK-X both achieved high computational efficiency and good performance in other studies and SPARK-X is the only method applicable to data with sample size exceeding 30,000 (Zhu et al. 2021). As expected, effect sizes are the major factor affecting performance. Larger effect sizes produced more distinct patterns, which were easier to be distinguished from random spatial distributions and resulted in higher AUC values. For hotspot and streak patterns, SpaGene, SpatialDE, and SPARK-X successfully distinguished spatially from non-spatially variable genes when patterns were distinct (AUC=1 at FC>=5 for hotspot and AUC=1 at FC>=8 for streak patterns). For less distinct patterns, SpaGene performed slightly better than SpatialDE and SPARK-X for smaller patterns, which obtained AUC of 0.64, 0.52 and 0.55 for SpaGene, SPARK-X and SpatialDE respectively at FC=2 and size=1 in hotspot patterns, while SPARK-X outperformed SpatialDE and SpaGene for bigger patterns (size>1) (Fig. 1C). For circularity and bi-quarter circularity patterns, SpaGene achieved much better performance than SpatialDE and SPARK-X. For the circularity pattern, SpaGene achieved AUC of 0.99 even for the smallest pattern at FC=3 and AUC of 1 at FC>=5. In comparison, SpatialDE only obtained AUC of 0.73 at FC=3, and SPARK-X failed to distinguish spatially from non-spatially variable genes even at FC=5 (AUC=0.5) for the smallest pattern (size=1). SpaGene and SpatialDE achieved AUC of 1 while SPARK-X only obtained AUC of 0.72 at FC=8 and size=1. Although the performance of SpatialDE and SPARK-X improved with increasing pattern sizes, SpaGene was more powerful than SpatialDE and SPARK-X (Fig. 1C). For the bi-quarter circularity pattern, SPARK-X failed even at the largest effect size for the two small patterns (AUC=0.5 at FC=10, size=1 or 2), while SpaGene achieved AUC>=0.9 and SpatialDE obtained AUC of 0.7-0.83 at FC>=3 for any pattern sizes (Fig. 1C). For the Purkinje layer pattern, SPARK-X failed at any effect sizes (AUC=0.5), while SpaGene achieved AUC of 0.81 at FC= 2, 0.99 at FC= 3 and 1 at FC>=5 (Fig. 1C). SpatialDE was not applied in this setting due to long computational time. To summarize, SpaGene achieved good performance for all spatial patterns, which obtained AUC>=0.98 at FC>=3 for relatively big patterns (size>1) and AUC close to 1 at FC>=5 for any pattern sizes. In comparison, SPARK-X seemed to be very sensitive to pattern shapes, which worked well for hotspot and streak patterns, but not for circularity, bi-quarter circularity and Purkinje layer patterns even when patterns were strongly distinct from the background. Furthermore, SpaGene was more robust against pattern sizes than SpatialDE and especially SPARK-X, which sometimes showed more power to identify indistinct and large patterns than small distinct patterns. For example, SPARK-X obtained AUC of 0.8 at FC=3 and size=3, but AUC of 0.7 even at FC=8 and size=1 for circularity patterns. SpatialDE obtained AUC of 0.7 at FC=3 and size=1, but 0.82 at FC=2 and size=5 for bi-quarter circularity patterns. We also simulated scenarios with varying number of genes and cells/locations (Fig. S1-S5). We found that the performance of SpaGene were less dependent on the number of cells/locations compared to SpatialDE and SPARK-X. The evaluation on the simulation datasets sampled from real data obtained similar results (Fig. S6-S9).

In terms of time complexity, SpaGene and SPARK-X are much more computationally efficient than SpatialDE. SpatialDE requires several orders of computational time than SpaGene and SPARK-X, and its runtime increases linearly or cubically with the number of genes and the number of cells/locations (Fig. S10A). For example, it takes SpatialDE 4,045 seconds to analyze a data with 10,000 genes and 5,000 cells/location, while it only takes SpaGene and SPARKX 11 and 22 seconds, respectively (Fig. S10B).

### Application to MOB by spatial transcriptomics

We applied SpaGene to spatial transcriptomics data from main olfactory bulb (MOB) (Stahl et al. 2016), involving 16,218 genes measured on 262 spots. The MOB has a roughly concentric arrangement of seven cell layers (Nagayama et al. 2014). SpaGene identified 634 as spatially variable genes (adjusted p-value, adjp<0.05), including genes known to be located in specific layers. Several examples were shown in Fig. 2A, such as *Pcp4* in Granule cell layer (GCL) (adjp=3e-6) (Sangameswaran et al. 1989), *Slc17a7* in Mitral cell layer (MCL) (adjp=7e-4) (Zhang et al. 2021), *Cck* in Glomerular layer (GL) (adjp=2e-3) (Sun et al. 2020b), *Serpine2* in External plexiform layer (EPL) (adjp=4e-3) (Mansuy et al. 1993) and *Fabp7* in Olfactory nerve layer (ONL) (adjp=4e-76) (Young et al. 2013). Based on those identified spatially variable genes, SpaGene successfully reconstructed the underlying seven-layered MOB structure (Fig. S11). To be noted, SpaGene identified a pattern corresponding to subependymal zone (SEZ) (pattern 4 in Fig. S11). SEZ was unidentifiable by spatially unaware single-cell clustering, which only discovered five distinct clusters (Fig. S12A). SEZ harbors neural stem cells. *Sp9* is the top gene specifically located in SEZ, which is a transcription factor that regulate MOB interneuron development (Li et al. 2018).

**Fig. 2.**
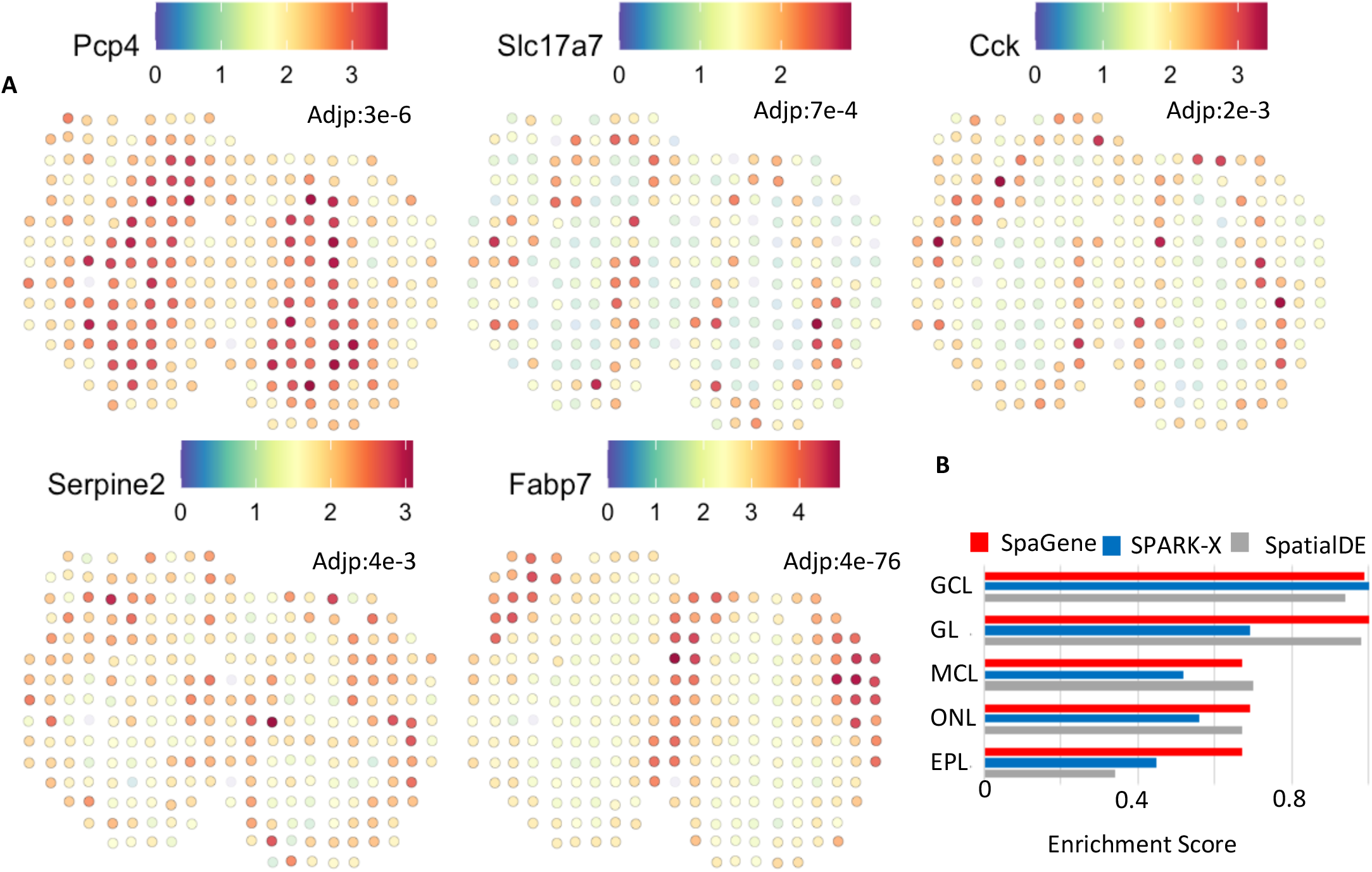
Application of SpaGene to spatial transcriptomics of main olfactory bulb data (MOB). A) Visualization of five known spatially variable genes located in specific MOB layers (high expression in red, and low in blue), with adjusted p-values rom SpaGene; B) Enrichment scores of markers in location-restricted cell types by SpaGene, SpatialDE and SPARK-X.

We compared SpaGene with SPARK-X and SpatialDE. Overall, SpaGene and SpatialDE had more overlapping than SPARK-X (Fig. S12B). We ranked spatially variable genes by each method and carefully examined those genes identified to be very significant by one method but insignificant by another method. First, we ranked genes by SpaGene and listed the top 6 genes with inconsistent results (Fig. S13). *Kif5b, Atf5, Sorbs1, Piekhb1* and *Mfap3l* were detected to be very significant by SpaGene (adjp<e-21), which were all specifically expressed in ONL (Fig. S11). However, none of them were found by SPARK-X, while *Atf5, Piekhb1* and *Mfap3l* were undiscovered by SpatialDE (Fig. S13). Another gene, *Grb2* was identified by SPARK-X but missed by SpatialDE, showing a very clear GCL pattern (Fig. S13). Then we ranked genes by SPARK-X and checked the top 6 inconsistent ones (Fig. S14). *Camk2a, Psd3, Meis2, Calm2, Arf3* and *Stxbp1* ranked high by SPARK-X, which displayed strong GCL patterns. All were identified by SpaGene but none by SpatialDE, indicating SpatialDE had limited power in identifying GCL-specific genes (Fig. S14). Finally, we ranked genes by SpatialDE and examined the top 6 inconsistent ones, including *Spem1, Siglec1, Cck, Kif5b, Apoe*, and *Il12a* (Fig. S15). *Spem1, Siglec1*, and *Il12a*, however, only expressed in one or two spots, which were likely to be false signals. *Cck, Kif5b* and *Apoe* exhibited GL or ONL patterns, which were identified by SpaGene but missed by SPARK-X (Fig. S15). These comparisons demonstrated that SpaGene successfully identified genes with visually distinct patterns, while SPARK-X and SpatialDE missed several genes in certain layers even they showed distinct patterns.

Since spatially unaware single-cell clustering uncovered cell types located in MOB layers, we expected that top markers in each layer-specific cell type would be identified as spatially variable genes. We calculated scores to measure the enrichment of those top markers in SpaGene, SPARK-X and SpatialDE. SpaGene obtained high enrichment scores in all layers, suggesting it successfully identified all layer-specific marker genes as being very significant. In contrast, SPARK-X obtained high in GCL layers but low in other layers. SpatialDE achieved high scores in Mitral cell layer, but relatively low scores in GCL and EPL layers (Fig. 2C).

### Application to mouse preoptic hypothalamus by MERFISH

We applied SpaGene to mouse preoptic hypothalamus data by MERFISH (Moffitt et al. 2018), consisting of 161 genes measured on 5,665 cells. The 161 genes include 156 pre-selected markers of distinct cell populations and five blank control genes. Spatially unaware single-cell clustering identified multiple cell types, most of which were spatially localized in specific regions, such as mature oligodendrocyte (OD), ependymal, mural and some inhibitory and excitatory neuron cell types (Fig. 3A). SpaGene identified those markers from region-specific cell types as top variable genes. Some representative genes were shown in Fig. 3B, such as *Ntng1* in inhibitory neurons (adjp=5e-108), *Mbp* in mature OD (adjp=0), *Cd24a* in Ependymal (adjp=0), *Adcyap1* in excitatory neurons (adjp=0), and *Myh11* in Mural cells (adjp=4e-24). Comparing SpaGene with SPARK-X and SpatialDE, we found their results were highly correlated in terms of significance (R=0.91 between SpaGene and SpatialDE, R=0.7 between SpaGene and SPARK-X, and R=0.8 between SPARK-X and SpatialDE) (Fig. 3C). We also compared the number of positive genes given the number of negative control genes identified (Fig. 3D). The results supported a higher power of SpaGene. For example, SpaGene detected 149 true positives, while SpatialDE discovered 144 and SPARKX revealed 128, when one negative control was detected (one false positive).

**Fig. 3.**
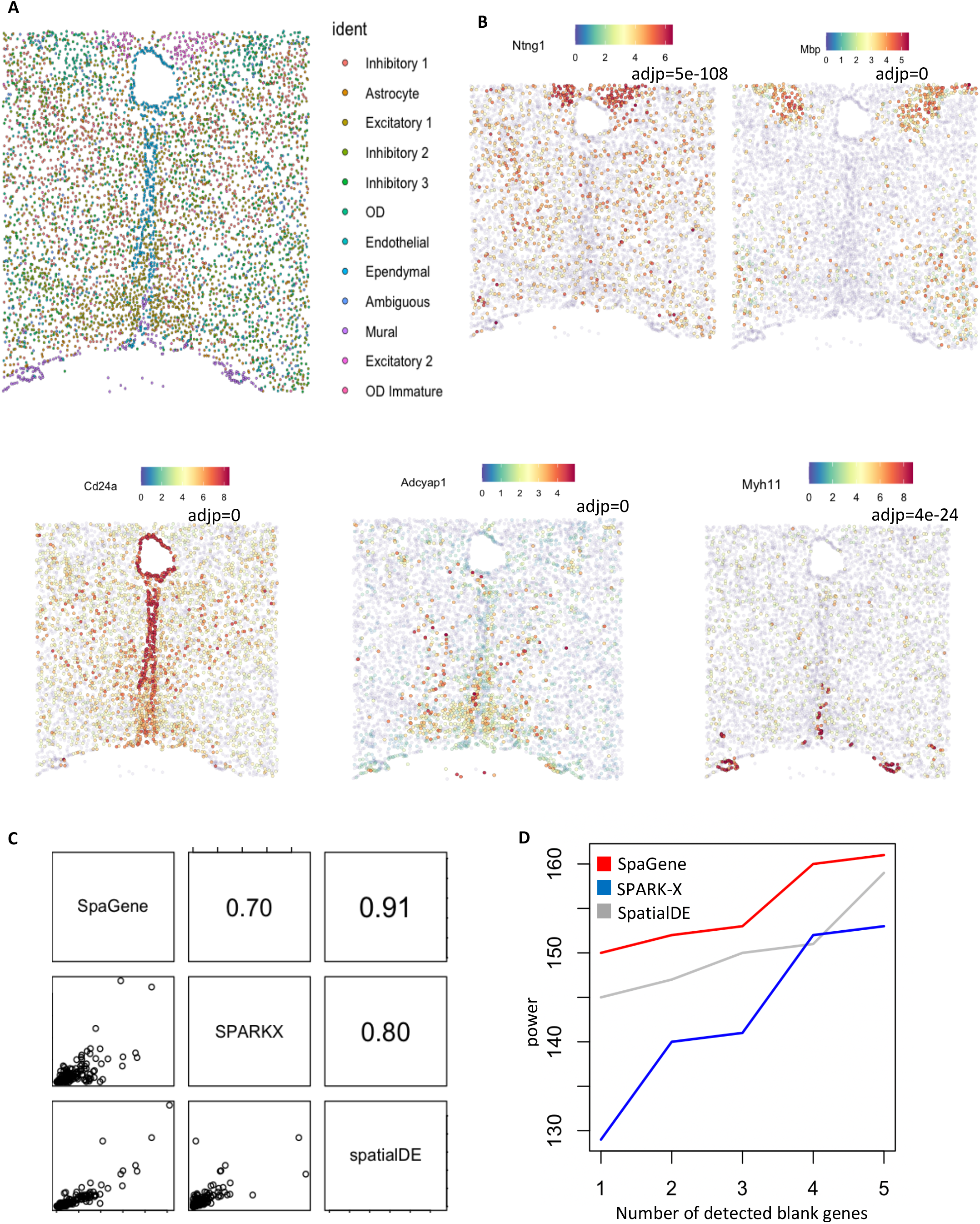
Application of SpaGene to MERFISH of mouse preoptic hypothalamus data. A) Spatially-unaware cell clustering; B) Visualization of five spatial variable genes (high expression in red and low in blue) with adjusted p-values from SpaGene; C) Pairwise correlation of results from SpaGene, SpatialDE and SPARK-X; D) Power plot shows the number of genes with spatial expression pattern (y axis) identified by SpaGene, SpatialDE and SPARK-X versus the number of blank control genes identified at the same threshold.

### Application to mouse cerebellum by Slideseq V2

We applied SpaGene to mouse cerebellum data by Slideseq V2 (Stickels et al. 2021), containing 20,141 genes measured on 11,626 spots. SpaGene identified 619 genes with spatial patterns (adjp<0.05). The cerebellum is made of three layers, molecular, Purkinje and granular layers from outer to inner, and white matter underneath. SpaGene detected genes, known to be specifically located in three layers and white matter, to be very significant, such as *Kcnd2* in granular layer (adjp=4e-253) (Varga et al. 2000), *Car8* in Purkinje layer (adjp=0) (Miterko et al. 2019), *Gad1* in molecular layer (adjp=2e-64) (Kirsch et al. 2012) and *Mbp* in white matter (adjp=0) (Verity and Campagnoni 1988) (Fig. 4A). Based on those identified spatially variable genes, SpaGene successfully reconstructed the tightly folded layer structure of cerebellum. Patterns 1 and 3 corresponded to granular layer, patterns 2, 6 and 8 represented molecular layer, patterns 4 and 5 stood for Bergmann glia and purkinje neurons in Purkinje layer, and pattern 7 imaged white matter (Fig. S16).

**Fig. 4.**
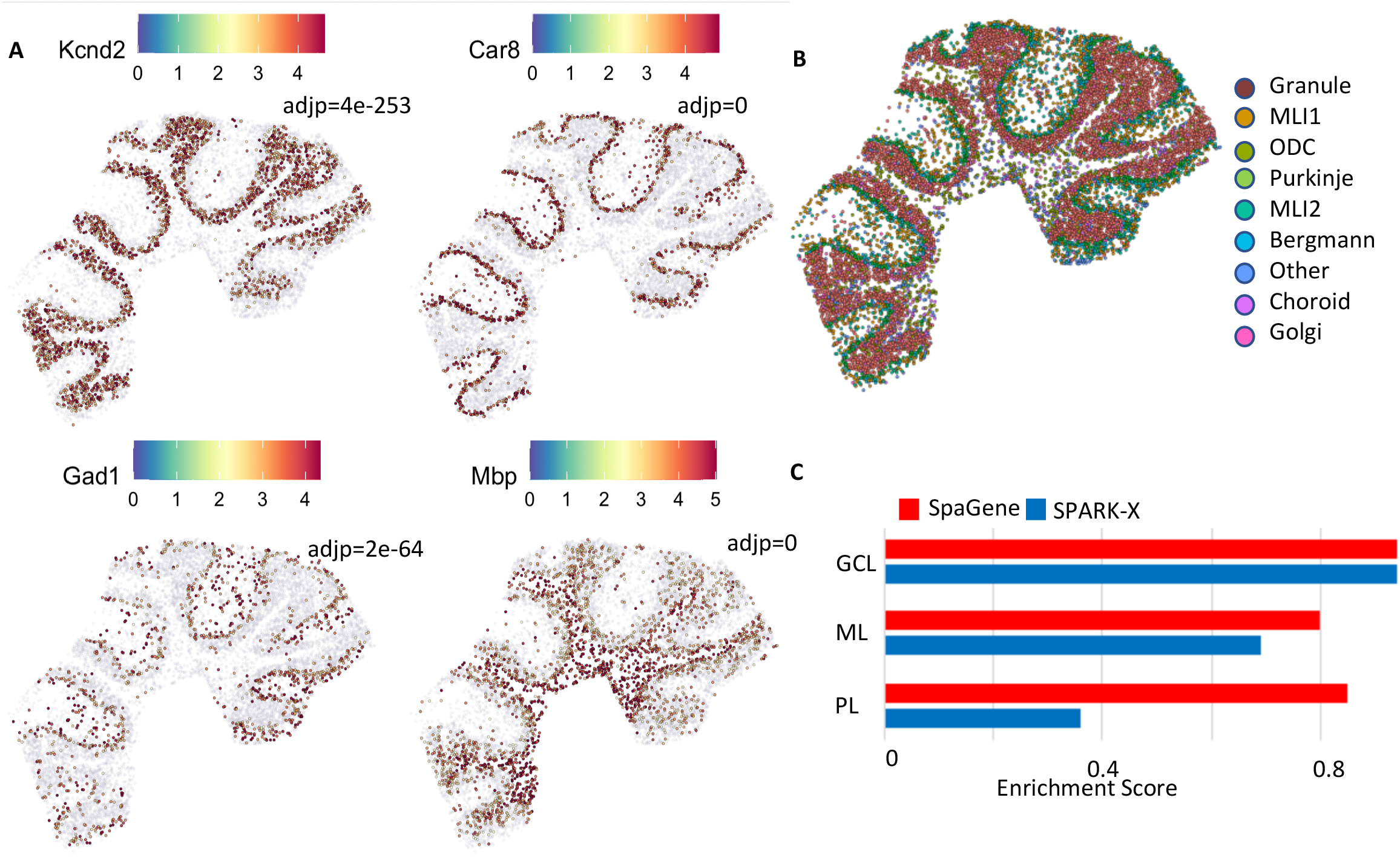
Application of SpaGene to Slideseq V2 of mouse cerebellum data. A) Visualization of four known spatially variable genes located in specific cerebellum layers (high expression in red, and low in blue), with adjusted p-values from SpaGene; B) Spatially unaware cell clustering; C) Enrichment scores of markers in location-restricted cell types by SpaGene and SPARK-X.

We compared SpaGene with SPARK-X but not SpatialDE because it would take hours to analyze such large-scale data. SPARK-X discovered 530 genes, while 230 overlapped with SpaGene (Fig. S17). We examined carefully at those genes detected to be very significant by one method but insignificant by the other one (Fig. S17). Those genes specifically located in Purkinje layer, such as *Car8, Itpr1, Pcp2*, and *Pcp4*, were detected as being the most significant by SpaGene (adjp=0) but undetected by SPARK-X, suggesting SPARK-X had limited power to identify the Purkinje pattern (Fig. S18). In comparison, *Catsperd, Ifit3*, and *Ptprt* ranked top by SPARK-X, but undetected by SpaGene, which didn’t seem to have obvious patterns (Fig. S19). SpaGene obtained the significance of *Mog* were just below the cutoff (adjp=0.05), which seemed to be dispersed in the white matter (Fig. S19).

Spatially unaware single-cell clustering found localized cell types, such as molecular layer neurons, purkinje neurons in the purkinje layer, granule cells in the granule layer (Fig. 4B). We expected markers in those spatially-restricted cell types were identified and ranked top by the methods. The enrichment analysis found that SpaGene obtained high enrichment scores in all three layers, while SPARK-X got a high score in granular layer, but low scores in other two layers, especially in the Purkinje layer. This result further demonstrated that SpaGene is more robust to any spatial patterns (Fig. 4C).

### Application to MOB by HDST

We applied SpaGene to olfactory bulb from high-definition spatial transcriptomics (HDST) (Vickovic et al. 2019), involving 19,950 genes measured on 181,367 spots. HDST is extremely sparse, where only 21 spots have more than 50 genes detected. In this case, SpaGene used an adaptive strategy to expand the neighborhood search for genes with high sparsity. SpaGene identified 249 genes as being spatially variable. The most significant genes included *Ptgds* (adjp=1e-232), *Gphn* (adjp=3e-114) and *Camk1d* (adjp=3e-61). Although spatial patterns of those genes were not visually distinct due to high sparsity of the HDST data, there were vague patterns showing *Ptgds* localized in ONL, *Gphn* in MCL and EPL, and *Camk1d* in GCL (Fig. 5A). Those specific localizations have been reported before (Rees et al. 2003; Perera et al. 2020) and validated by in situ hybridization in the Allen Brain Atlas (Fig. 5B).

**Fig. 5.**
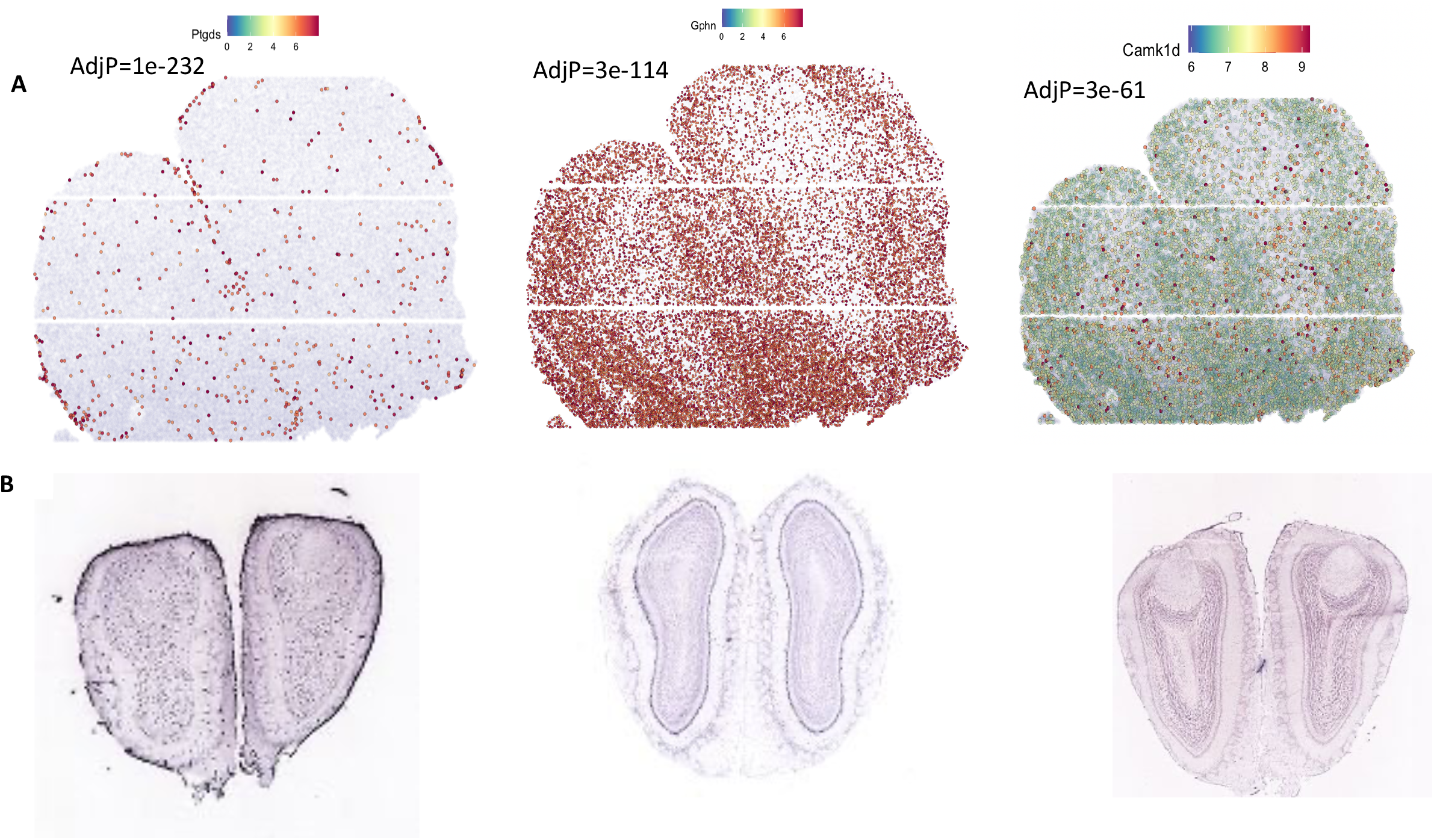
Application of SpaGene to HDST of MOB data. Visualization of three spatially variable genes. A) gene-expression levels from HDST (high in red, low in blue), with adjusted p-values from SpaGene; B) in situ hybridization results for the three genes obtained from the Allen Brain Atlas.

We compared SpaGene with SPARK-X but not SpatialDE because it would take months to analyze such large-scale data. SPARK-X detected 133 genes, which overlapped significantly with SpaGene (90 in common). Among the 40 genes most associated with each MOB layer (top 5 genes in eight patterns in Fig. S11), SpaGene found 12 genes (*Ptgds, Fabp7, Gad1, Vtn, Kctd12, Kif5b, Apod, Pcp4, Gpsm1, Slc1a2, Nrgn, and Map1b*), while SPARK-X only detected six (*Ptgds, Fabp7, Kctd12, Kif5b, Apod*, and *Pcp4*).

### Identification of spatially colocalized ligand-receptor pairs

We extended SpaGene to identify cell-cell communications mediated by colocalized ligand and receptor pairs. SpaGene found 35 ligand-receptor interactions from the mob data by spatial transcriptomics. The two most significant ligand-receptor pairs were Igfbp5-Cav1 (adjp=3e-31) and Apoe-Lrp6 (adjp=1e-18), both happening between ONL and GL. *Apoe* is known to be enriched in ONL and GL and also identified to be very significant by SpaGene (adjp=1e-50). Most spots with high *Apoe* expression were surrounded with spots with high *Lrp6* expression (Fig.6A), suggesting potential interactions between them. Apoe-Lrp6 mediates Wnt signaling, which is important for the regulation of synaptic integrity and cognition (Zhao et al. 2018). The identification of Apoe-Lrp6 between ONL and GL layers might be suggestive of the potential regulation of Wnt signaling in the establishment of periphery–CNS olfactory connections.

SpaGene found 13 ligand-receptor interactions from the mouse cerebellum data by Slideseq V2. The most significant pair was Psap-Gpr37l1 (adjp=1e-27) (Fig. 6B). Gpr37l1 was known to be strongly expressed in Purkinje layer and also identified by SpaGene (adjp=8e-130). *Psap*, in contrast, was not as specifically localized as *Gpr37l1* (adjp=6e-8). Psap-Gpr37l1 protects neural cells from cellular damage (Li et al. 2017). The identification of Psap-Gpr37l1 between Purkinje layer and surrounding layers further supports its important role in brain function. Additionally, Ptn-Ptprz1, identified as the only interaction by MERINGUE (Miller et al. 2021), ranked the top four by SpaGene (adjp=2e-7).

**Fig. 6.**
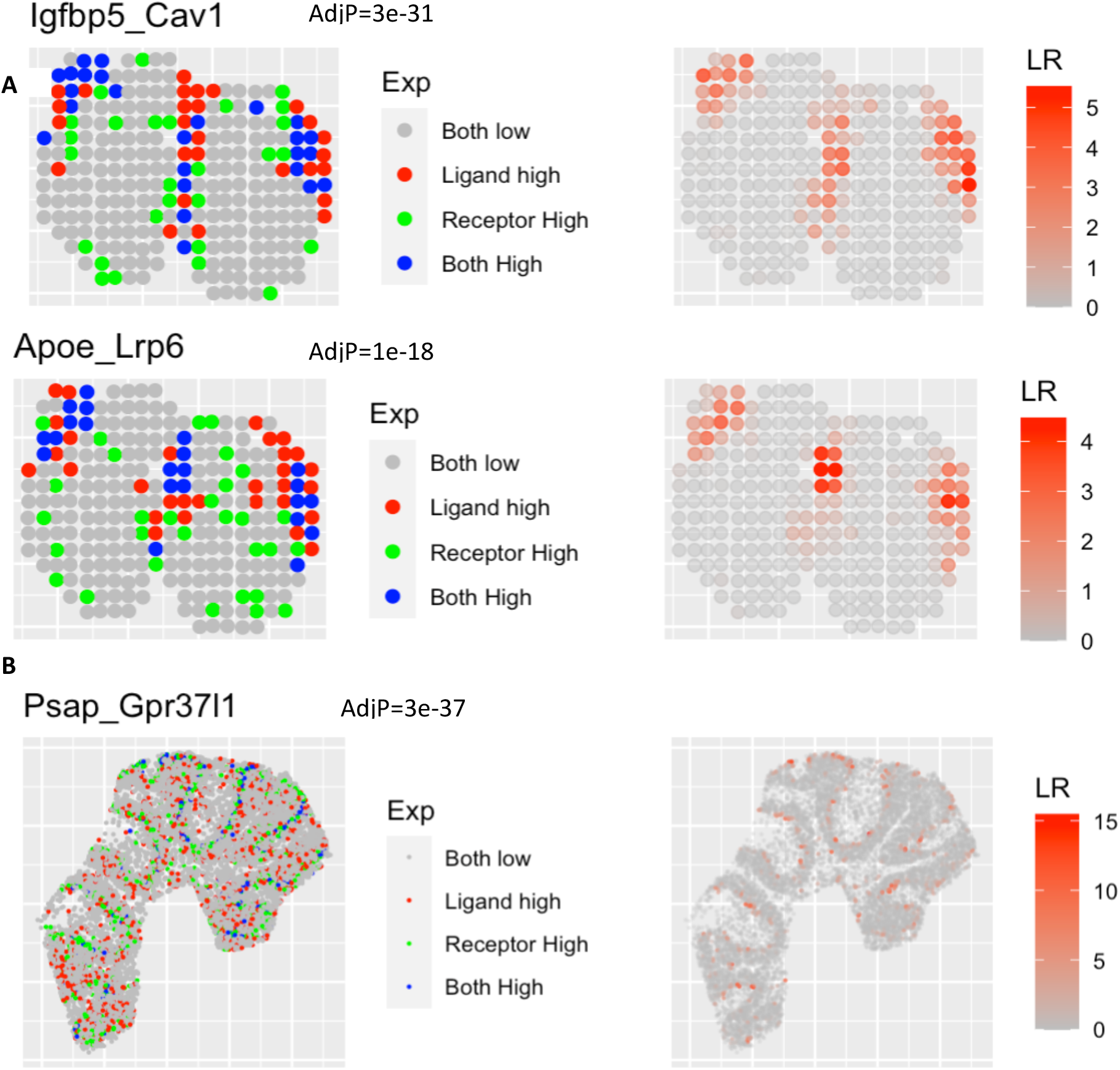
Extension of SpaGene to identify ligand-receptor interactions. A) Visualization of Igfbp5-Cav1 and Apoe-Lrp6 interactions for ST MOB data, with adjusted p-values from SpaGene. B) Visualization of the Psap-Gpr37l1 interaction for Slideseq V2 mouse cerebellum data, with the adjusted p-value from SpaGene. Left is the relative expression of the ligand and the receptor, right is the interaction strength.

## DISCUSSION

Recent advances in spatial omics technologies increase the demand for scalable and robust methods to characterize spatially variable patterns. Here, we developed SpaGene, a fast and model-free method to identify spatially variable genes. SpaGene has been extensively evaluated on seven datasets generated from a variety of spatial technologies, ranging from low to high throughput and spatial resolution. Additional analyses on breast cancer from spatial transcriptomics, mouse brain from 10X Visium, and olfactory bulb from Slide-seqV2 were shown in Supplementary Figures S20-S30. The results consistently demonstrated that SpaGene successfully identified known spatially variable genes and also markers in spatially-restricted cell clusters. Simple factor analysis on those identified genes reconstructed underlying tissue structures, further demonstrating the ability of SpaGene to characterizing spatial patterns.

Compared with existing approaches, SpaGene is more robust to pattern shapes, data distribution and sparsity, non-uniform cellular densities, and the number of spatial locations. The power of SpatialDE, SPARK and SPARK-X highly depend on spatial covariance models, that is, how well those predefined kernel functions match the true underlying spatial patterns. Moreover, SpatialDE and SPARK use parametric modeling based on the assumption of spatial data following Gaussian or Poisson distributions. Therefore, their performance would be compromised significantly for those genes whose expression misalign the model defined by those kernel functions and whose distribution violate Gaussian or Poisson distributions. SpaGene, in contrast, is a model-free and distribution-free method. Without any assumption, SpaGene is able to identify any spatial patterns and applied on any spatial omics data, such as identification of spatially localized clones and histone markers in spatial genomics and epigenomics data. The significance from SpaGene reflects the distinctness of spatial patterns rather than the extent of match to the defined model. SpaGene uses neighborhood graphs to represent spatial connections, making it more robust to non-uniform cellular densities common in tissues. Furthermore, SpaGene is highly computationally efficient. It only took seconds to minutes for SpaGene to analyze large-scale spatial transcriptomics data, which required hours, days or even months for most methods (Zhu et al. 2021) (Fig. S10C).

SpaGene is very flexible, which can tune neighborhood search spaces automatically based on the data sparsity. SpaGene can incorporate the cell type information to find spatially variable genes within the same cell type. For example, SpaGene identified *Aldoc* as the most spatially variable genes within the Purkinje layer (adjp=4e-90) (the function SpaGene_CT was provided in the package), which has been demonstrated to show a regional enrichment pattern that was consistent with the known paths of parasagittal stripes across individual lobules (Kozareva et al. 2021). Furthermore, SpaGene was easily extended to find colocalized gene pairs. It successfully identified Psap-Gpr37l1 and Ptn-Ptprz1 in mouse cerebellum, and Fn1-Cd44 in invasive breast cancer regions (Fig. S30). The default neighborhood search regions could be further adjusted to identify those long-distance interactions. In summary, SpaGene is very powerful tool to characterize any localized and co-localized patterns. Potential extensions of SpaGene to find alterations in spatial patterns across conditions would further expands its application.

## METHODS

### Method overview

Spatially variable genes are those with uneven spatial distribution of expression, where cells/spots with high expression are more likely to be spatially connected than random. Given a set of spatial locations, SpaGene first builds the spatial network using k-nearest neighbors. SpaGene then quantifies the spatial connection of cells/spots with high expression by their degree distribution. Finally SpaGene compares the observed spatial connection with those from random permutations (Fig. 1A). Genes with significantly higher spatial connection than random are identified as spatially variable genes.

The degree distribution p(i) is defined to be the fraction of cells/spots with degree of i. Earth mover’s distance (*EMD*) is used to quantify the distance from the observed degree distribution to a distribution from a fully connected network. Smaller EMD distances indicate higher spatial connection.

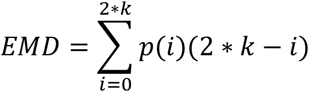

To generate the null distribution of *EMD*, the same number of cells/spots is randomly sampled and the spatial connection of those cells/spots is quantified as EMD’. The mean and the standard deviation of EMD’ is estimated after 5,00 random permutations. The observed EMD is compared to the null distribution of EMD’ to evaluate its significance.

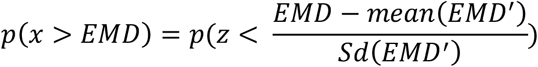

### Identification of spatial patterns

Non-negative matrix factorization is applied on spatially variable genes detected by SpaGene to identify distinct spatial patterns. NMF is implemented by the RcppML R package. The Spearman correlation between expression of spatially variable genes and cells/spots factor matrix from NMF is used to find the most representative genes in each pattern.

### Simulation designs

We followed simulation designs of SPARK-X and Trendsceek. Briefly we generated two datasets with five spatial expression patterns, local hotspot, streak, circularity, bi-quarter circularity and mouse purkinje layer. For the first four patterns, spatial locations of cells were generated by a random-point-pattern Poisson process. The spatial locations of the pattern of mouse purkinje layer was obtained from Slideseq V2 mouse cerebellum data. The expression values were either generated from negative binomial distributions following SPARK-X or bootstrap-sampled from spatial transcriptomics MOB data following Trendsceek. Simulation datasets varied on a number of parameters: 1) the number of genes varied from 1000, 3000, and 10,000, among of which 500 genes are spatially variable; 2) the number of cells varied from 300, 1000, 2000 and 5000 except for the purkinje layer pattern; 3) the fold change of expression in the spatial region compared to those in the background; For the negative binomial distribution, the fold change varied from 2, 3,5, 8 to 10. For the resampled real dataset, the expression of spiked cells were generated from 65%, 70%, 80% to 90% quantile of the expression distribution; 4) the number of spiked cells except for the purkinje layer pattern. For the hotspot and the streak patterns, the percentage of spiked cells varied from 5%, 10%, 20% to 30%. For the circularity and bi-quarter circularity patterns, the width of circularity varied between 0.05, 0.075, 0.1, 0.125 and 0.15.

### Spatial transcriptomics datasets

SpaGene was applied on seven spatial transcriptomics datasets, covering a variety of platforms with low and high throughput and spatial resolution. Two spatial transcriptomics data from mouse olfactory bulb and human breast cancer contained genome-wide expression profiles on only hundreds of spots (low spatial resolution) (Stahl et al. 2016). MERFISH on the mouse preoptic region of the hypothalamus targeted only 160 genes at single cell resolution. 10X Visium on the mouse brain comprised of whole transcriptomics on thousands of spots with a spatial resolution of 55 µm. Two Slideseq V2 from mouse cerebellum and olfactory bulb contained whole transcriptomics on tens of thousands of spots with a spatial resolution of 10 um. HDST from mouse olfactory bulb measured whole transcriptomics on hundreds of thousands of spots with a spatial resolution of 2µm.

## DATA ACCESS

Seven spatial transcriptomics data were available from original studies and also from the SpaGene Github repository https://github.com/liuqivandy/SpaGene. In addition to identification of spatially variable genes and patterns, the R package SpaGene also provides functions to visualize spatial patterns and co-localized ligand-receptor pairs. Vignettes on seven spatial transcriptomics data with raw data, codes and results, including spatial variable genes identification, pattern identification and visualization, co-localized ligand-receptor pairs identification and visualization, are also available at the GitHub.

## ACKNOWLEDGMENTS

This work is supported by National Cancer Institute grants (U2C CA233291 and U54 CA217450), and Cancer Center Support Grant (P30CA068485).

